# *Mycobacterium tuberculosis* effector protein PE5 hijacks the host CRL2 ubiquitin ligase complex

**DOI:** 10.1101/2025.06.11.659056

**Authors:** Bala T.S.A. Madduri, Omair Vehra, Ahri Han, Joycelyn Radeny, Carlos Resstel, Samantha L. Bell

## Abstract

*Mycobacterium tuberculosis* (Mtb) is the leading infectious killer and infects one quarter of the global population. During infection, Mtb evade host immune responses via secreted effector proteins that interfere with, modulate, and protect from potent antibacterial responses. One such class of mycobacterial effector proteins are the PE/PPE proteins, a huge family of proteins encoded by an impressive 10% of the Mtb genome. Because some PE/PPEs have demonstrated roles in immune regulation and host cell interaction, and because of the sheer number of PE/PPEs in pathogenic mycobacteria (∼170-200+ members depending on the species), they are thought to contribute to Mtb virulence and pathogenesis. However, the cellular functions of Mtb’s 169 PE/PPE proteins have yet to be comprehensively characterized, at least in part because of their high GC content and large regions of extremely repetitive regions found both in DNA and amino acid sequences. One member of this family, PE5, is likely critical for mycobacterial pathogenesis, but its cellular and molecular functions, particularly within a host cell, are unexplored. We investigated the molecular functions of PE5 using affinity purification coupled mass spectrometry (AP-MS) and identified an interaction with the CRL2 complex, a host E3 ubiquitin ligase complex. PE5 interacts with CRL2 through its C-terminal Gly-Gly motif, which is bound by the CRL2 substrate receptor, KLHDC2. PE5 does not get ubiquitinated, but it is degraded upon being bound by KLHDC2. Interestingly, binding to PE5 increases the autoubiquitination of KLHDC2, but this autoubiquitination does not interfere with KLHDC2’s ability to degrade known substrates or the ability of CRL2 to degrade substrates bound by other substrate receptors. Therefore, while PE5 binds to CRL2-KLHDC2 and does not get ubiquitinated itself, it does not appear to impede CRL2 activity. This study identifies a novel interaction between PE5 and a host ubiquitin ligase pathway, and it raises new questions about how PE5’s interaction with the host modulates cell biology to promote Mtb virulence. Furthermore, it implicates CRL2 complexes in mediating cell-intrinsic host response to Mtb infection, a novel function for this ubiquitin ligase complex. Our results expand our understanding of how PE/PPEs may target innate immune responses and contribute to our knowledge of how uncharacterized PE/PPEs contribute to Mtb’s virulence.

## INTRODUCTION

Tuberculosis (TB) is an ancient disease that remains the leading cause of death from an infectious agent. It is caused by *Mycobacterium tuberculosis* (Mtb), which infects one-quarter of the global population and kills more than 1.25 million people annually^1^. Despite the prevalence of this infection, we still lack a complete understanding of how Mtb evades host immune responses to cause disease, hampering efforts to eradicate TB.

Mtb infection begins with a macrophage phagocytosing Mtb, and it progresses as a tug-of-war between a macrophage’s potent antibacterial responses and Mtb’s numerous immune evasion strategies^2^. These initial host-pathogen interactions critically influence disease progression^3^. Key to Mtb’s innate immune evasion are its many secreted effectors that interfere with and modulate host responses^4^. One major class of mycobacterial effector proteins are the PE/PPEs, which are encoded by an impressive 10% of the Mtb genome and contains ∼169 members in Mtb^5–7^. This family of effectors is named after the conserved Pro-Glu (PE) or Pro-Pro-Glu (PPE) residues in their N-terminal domains. PE/PPEs’ C-terminal domains are highly variable with diverse structures and functions, many of which are disordered or otherwise uncharacterized. Despite the widely held assumption that this large family of effectors is important for Mtb virulence, only a few members have been studied in detail. Indeed, several PE/PPEs have been shown to interfere with host responses to Mtb infection^8^. However, the large size of this effector family of effectors, coupled with the challenges of studying many members (high GC content (up to 80-85%), repetitive sequences (both DNA and amino acid), and seeming redundancy), has left the PE/PPEs understudied and their molecular functions during infection poorly understood.

PE5 is one member of the PE/PPE family that has been studied in the context of bacterial physiology but has poorly understood functions mediating host-pathogen interactions. Like many PEs, PE5 is encoded in an operon with its PPE pair, PPE4, and PE5/PPE4 is part of the virulence-associated ESX-3 secretion system, one of Mtb’s 5 type VII secretion systems^9^. These two PE/PPEs form a heterodimer for secretion via ESX-3, and their secretion is critical for iron acquisition; therefore, they are essential in Mtb. As a result, knockouts can only be generated when iron is supplemented in the form of heme. However, the specific role of PE5/PPE4 in iron acquisition remains unclear. Previous reports have demonstrated that PE5, but not PPE4, is robustly secreted by Mtb into culture filtrates^9–11^, suggesting PE5 is a secreted effector with a function outside of the bacterium. In addition, it is highly conserved among pathogenic mycobacteria, which provides evolutionary evidence of its importance in mycobacterial virulence^12–15^.

Cullin RING ubiquitin ligases (CRLs) are a large family of modular E3 ubiquitin ligase complexes that use interchangeable receptors to identify substrates for ubiquitination. Cullin-2-containing complexes, or CRL2 complexes, are equipped with substrate receptors that recognize specific C-terminal motifs called ‘C-degrons’^16^. C-degrons typically arise from proteolysis or translation defects, so CRL2 mediates the ubiquitination of C-degron-containing proteins to facilitate their proteasomal degradation^17^. CRL2 uses a panel of substrate receptors that each recognize different C-degrons that are less abundant in the standard proteome, which biases CRL2 complexes to degrade damaged or defective proteins. However, many proteins naturally contain C-degrons, and it remains unclear how CRL2 complexes might affect the stability of normal proteins in the cell. CRL2 influences multiple cellular processes^18,19^, but its regulation and function in immune cells, and therefore, its role during Mtb infection, are unknown.

Affinity purification coupled mass spectrometry (AP-MS) is a powerful tool to identify novel protein-protein interactions and probe the complex interplay between pathogens and host cells. AP-MS has been especially successful in describing uncharacterized Mtb effectors, and several previous studies have successfully used this approach to gain valuable insight into Mtb-macrophage interactions. One recent study identified 187 Mtb-human protein-protein interactions involving 34 Mtb secreted effectors^3,20^, and found an interaction between LpqN and the host ubiquitin ligase CBL, which reprograms macrophages toward antiviral signaling at the expense of antibacterial responses^20,21^. Another study ectopically expressed PE_PGRS47 in macrophages and found it interacted with Rab1A, resulting in blunted autophagic induction during Mtb infection^22^. Similarly, a PE_PGRS protein, MirA from the Mtb relative *M. marinum* was found to interact with the host actin nucleator N-WASP to induce actin polymerization and form actin tails that enable cell-to-cell spread^23^.

Here, we used AP-MS to discover that in mammalian cells, PE5 interacts with CRL2 complexes via the substrate receptor KLHDC2, which recognizes the C-degron found in PE5. Our studies demonstrated a specific interaction between PE5 and CRL2-KLHDC2, but PE5’s lack of ubiquitination by CRL2 led us to hypothesize alternative scenarios beyond PE5 being a substrate of this complex. We explored a potential role for PE5 as an inhibitor of CRL2 complexes but found no change in the activity of the CRL2 complexes we examined. Nonetheless, the substrate receptor KLHDC2 is required to control intracellular bacterial replication, suggesting it has an antibacterial role during infection. Likewise, ectopic expression of PE5 promotes intracellular bacterial replication, suggesting it blocks host responses. PE5’s ability to promote infection and bind to a host restriction factor supports a role for PE5 as a secreted effector protein that targets and blocks antibacterial host responses.

## RESULTS

### PE5 binds to KLHDC2, a substrate receptor of the CRL2 ubiquitin ligase complex

To gain insight into PE5’s cellular and molecular functions during infection, we explored its protein-protein interactions in a mammalian cell. To do this, we used a well-established unbiased approach of affinity purification coupled with mass spectrometry (AP-MS), which has been used previously to identify novel functions of Mtb secreted effector proteins^20^. We cloned the sequence of PE5 from Mtb H37Rv into a mammalian expression vector and inserted an N-terminal GFP tag. We transiently transfected this GFP-PE5 plasmid into HEK293T cells for ectopic expression of PE5, pulled down PE5 using GFP-trap resin, and identified PE5’s protein-protein interactions via mass spectrometry (Figure 1A). Through this approach, we identified multiple host proteins enriched in the pulldown of GFP-PE5 compared to GFP alone. Among these, we identified several members of the CRL2 complex (Figure 1B). CRL2 is a multi-subunit ubiquitin ligase complex^17^ where Cullin-2 (CUL2) recruits adapters Elongin B and C, which then recruit one of 40+ interchangeable substrate receptors^16^. These receptors recognize unique C-degrons to target proteins for proteasomal degradation^16,17^ (Figure 1C). Among the interactors we identified for PE5 (Figure 1B-C) was the substrate receptor KLHDC2, which recognizes the C-degron Gly-Gly (GG) in substrates^24^. This is consistent with PE5’s C-terminal Gly-Gly motif (Figure S1A). We validated the identified AP-MS interactions between PE5 and members of the CRL2 complex using directed co-immunoprecipitations (co-IPs) with GFP-PE5 and 3xFLAG-tagged members of the CRL2 complex (Figure 1D-E). While PE5 strongly interacted with KLHDC2, it bound less robustly to other members of the CRL2 complex, suggesting that these co-IPs failed to capture the full assembled complex or that PE5 binds primarily to free KLHDC2 rather than CRL2-KLHDC2 complexes.

**Figure 1.**
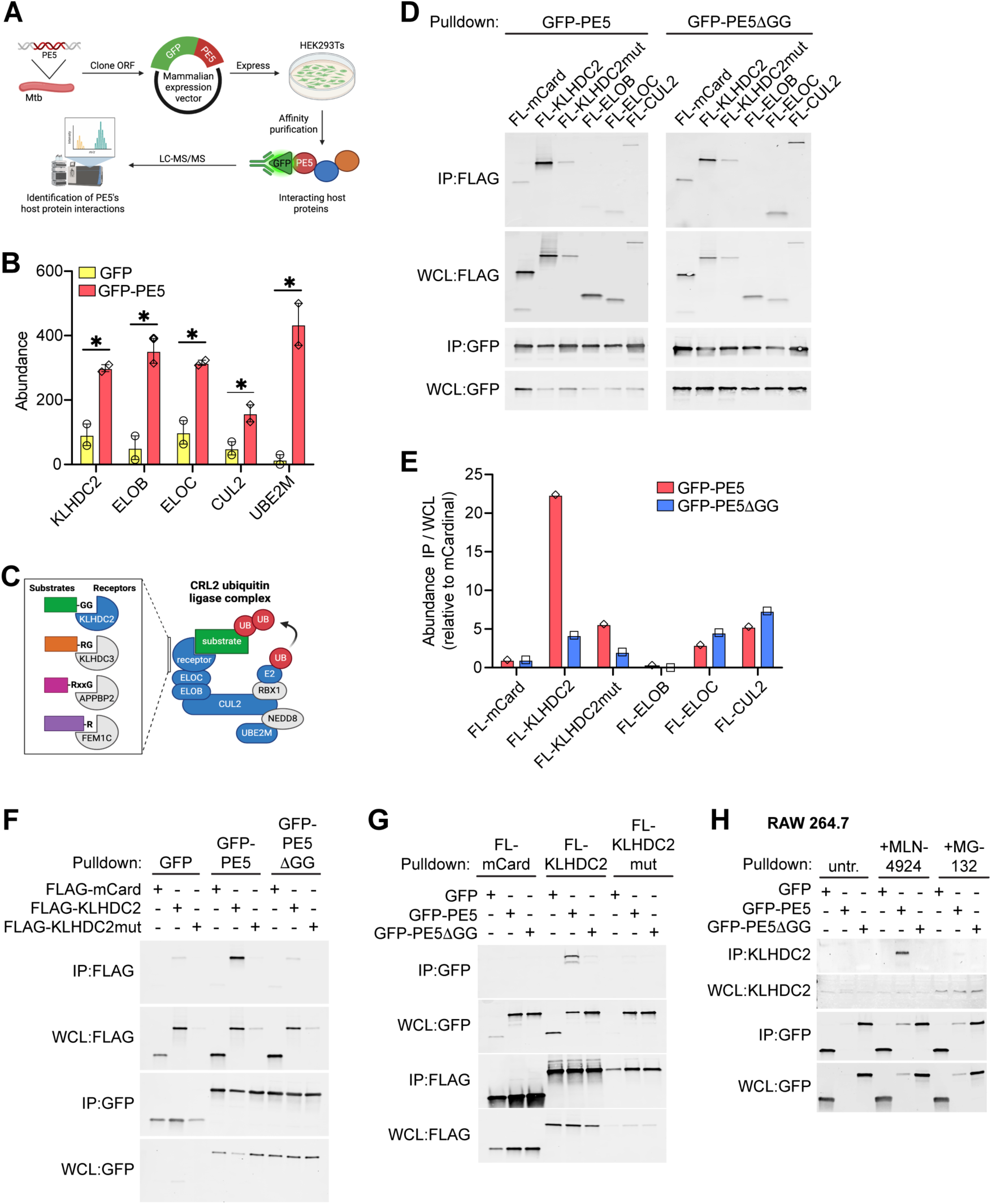
PE5 binds to KLHDC2, a substrate receptor of CRL2. (A) Schematic of the experimental strategy of AP-MS generated with BioRender. N-terminal-GFP tagged-PE5 or GFP were transiently expressed in HEK293T cells for 48 hours, and GFP-trap mediated AP-MS was used to identify protein-protein interactions. (B) Abundances of CRL2 complex member identified by mass spectrometry from HEK293Ts cells expressing GFP-PE5 or GFP alone (n=2). (C) Schematic generated with BioRender of the CRL2 complex, substrate receptors, and typical C-degrons recognized by each. (D) Co-IP (GFP-trap) from HEK293Ts transiently transfected with GFP-PE5 or GFP-PE5ΔGG and a panel of 3xFLAG (FL)-tagged CRL2 complex members identified by AP-MS or mCardinal as a negative control. Whole cell lysates (WCL) and immunoprecipitations (IP) shown for both bait (GFP-tagged proteins) and prey (FL-tagged proteins). (E) Quantification of D expressed as the fold enrichment of a protein in the IP relative to the WCL. Values were then normalized to the negative control mCardinal (mCard, set to 1). (F) Co-IP (GFP-trap) of RAW 264.7 cells stably expressing GFP (negative control), GFP-PE5, or GFP-PE5ΔGG, along with 3xFLAG (FL)-tagged mCardinal (negative control), KLHDC2, or KLHDC2 S269E (KLHDC2mut). (G) Similar to F but a co-IP using FLAG resin, making FL-tagged proteins the bait and GFP-tagged proteins the prey. (H) Co-IP (GFP-trap) from RAW 264.7 cells stably expressing GFP, GFP-PE5, or GFP-PE5ΔGG and probed for endogenous KLHDC2. Cells were treated with MLN4924 (5 mM) or MG-132 (20 mM) for 2 hr as indicated to increase PE5 expression to detectable levels. Results in D-H are representative of at least three independent experiments. *p < 0.05 by two-tailed t-test. Error bars indicate SD.

PE5’s C-terminal Gly-Gly motif and KLHDC2’s recognition of a Gly-Gly C-degron suggest a clear mechanism for this protein-protein interaction. To test this, we generated a mutant of PE5 lacking its C-terminal Gly-Gly residues (PE5ΔGG) and performed directed co-IPs with CRL2 complex members. While PE5 interacts robustly with KLHDC2 (Figure 1D-E), the PE5ΔGG mutant does not interact with KLHDC2 (Figure 1D-E). Next, to investigate how KLHDC2 binds PE5, we generated a variant of KLHDC2 with a mutation in its substrate binding domain (S269E, ‘KLHDC2mut’) and found that this KLHDC2 mutant could no longer bind PE5 or PE5ΔGG (Figure 1D-E). Together, this indicates that KLHDC2 binds the C-terminal Gly-Gly in PE5 via its substrate-binding domain.

Because we performed AP-MS and directed co-IPs via transient transfection in HEK293T cells in order to facilitate high levels of protein production, we next sought to validate the PE5-KLHDC2 interaction in macrophages, the cell type most relevant for Mtb infection. To do this, we stably expressed GFP, GFP-PE5, and GFP-PE5ΔGG in combination with 3xFLAG-mCardinal (negative control far-red fluorescent protein), 3xFLAG-KLHDC2, and 3xFLAG-KLHDC2mut in RAW 264.7 macrophages using lentiviral transduction and drug selection. Using these cell lines, we pulled down GFP-PE5 and, in line with our findings in HEK293Ts, PE5 interacted with 3xFLAG-KLHDC2 in macrophages (Figure 1F). We also performed the reverse co-IP, pulling down 3xFLAG KLHDC2 and probing for GFP-PE5, and once again found a strong interaction (Figure 1G). Finally, to validate that this interaction is physiologically relevant, we assessed the interaction between ectopically expressed PE5 and endogenous KLHDC2. To increase PE5 protein levels in RAW 264.7 cells, we treated cells with either the pan-cullin inhibitor (MLN-4924) or the proteasomal inhibitor (MG-132), both of which effectively stabilized PE5 (see below), prior to performing co-IPs. Here again, we found that PE5, but not PE5ΔGG, interacted with endogenous KLHDC2 in macrophages (Figure 1H).

### The PE5-KLHDC2 interaction is highly specific

Having established a strong interaction between PE5 and KLHDC2, we next sought to further explore the specificity of this interaction. Although other CRL2 substrate receptors were not identified by AP-MS, we performed directed co-IPs to test whether other substrate receptors, which recognize C-degrons not found in PE5, could nonetheless bind to PE5. While KLHDC2 recognizes a C-terminal Gly-Gly^24^, KLHDC3 recognizes a C-terminal Arg-X-Arg/Lys/Gln-Gly^25^; FEM1B/C recognizes a C-terminal Arg, and APPBP2 recognizes a C-terminal Arg-X-X-Gly (Figure 1C)^16,17,26^. None of these substrate receptors were pulled down with PE5 (Figure 2A-B).

**Figure 2.**
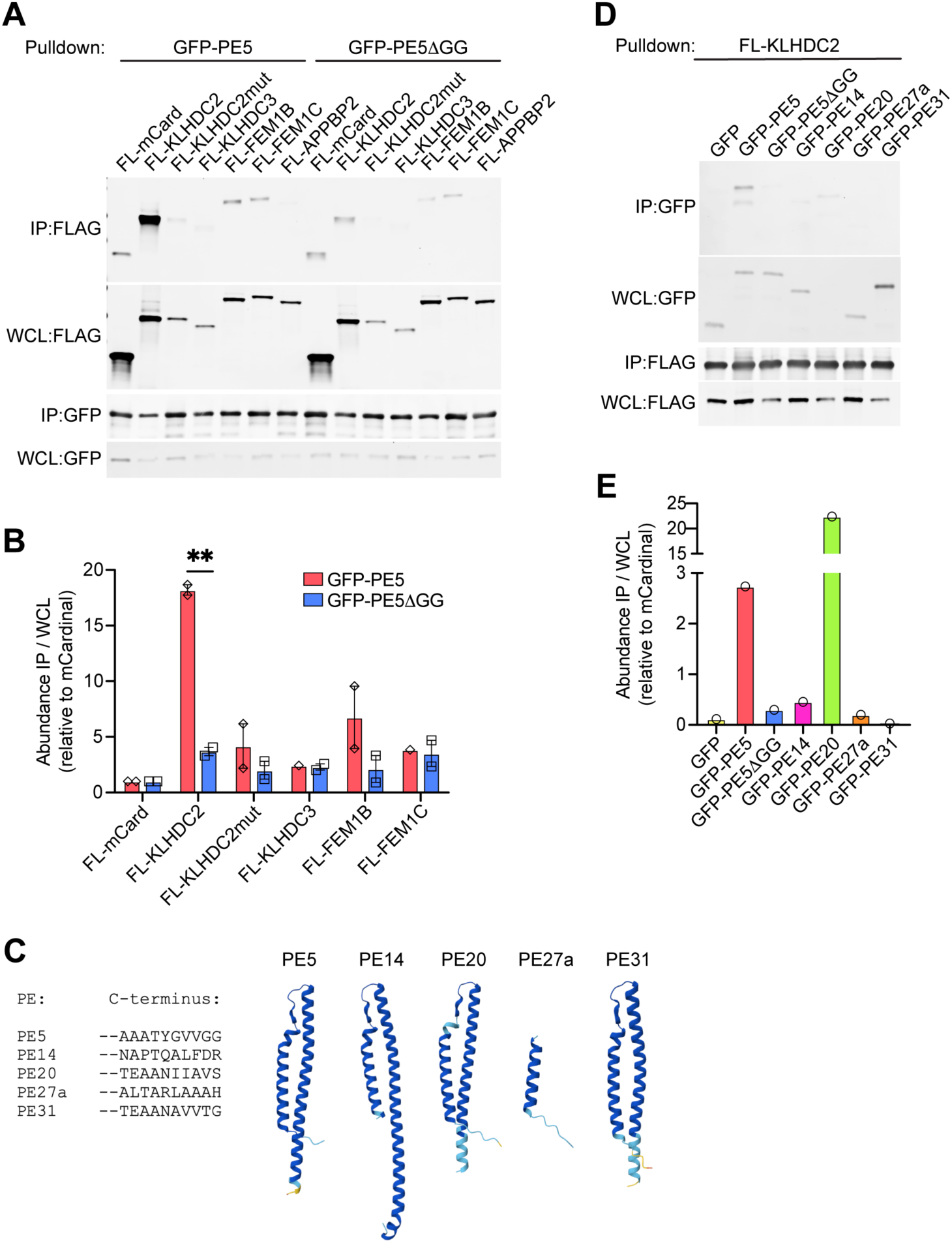
PE5 and KLHDC2 specifically interact. (A) Co-IP (GFP-trap) from HEK293Ts transiently transfected with GFP-PE5 or GFP-PE5ΔGG and with a panel of 3xFLAG (FL)-tagged CRL2 substrate adaptors, including PE5’s identified binding partner, KLHDC2 and KLHDC2 S269E (mut). (B) Quantification of A expressed as the fold enrichment of a protein in the IP relative to the WCL (n=2). Values were then normalized to the negative control mCardinal (mCard, set to 1). (C) The 10 C-terminal amino acids from a panel of tested PE proteins (left), and AlphaFold predicted structures of each PE protein (right). (D) Co-IP (FLAG resin) from HEK293T transiently transfected with 3xFLAG-tagged KLHDC2 and a panel of GFP-tagged small PEs (PE5, PE5ΔGG, PE14, PE20, PE27a, PE31). (E) Quantification of D expressed as the fold enrichment of a protein in the IP relative to the WCL. Data in A-B and D-E are representative of at least three independent experiments. **p < 0.005 by two-tailed t-test. Error bars indicate SD.

We next tested the possibility that KLHDC2 could bind nonspecifically to all PEs. To test this, we cloned several additional PEs from Mtb (PE14, PE20, PE27a, PE31), adding an N-terminal GFP tag and inserting them into mammalian expression vectors. We chose PEs that were similar in size to PE5, containing the PE domain (2 alpha helices) and flexible C-terminal tails (Figure 2C). Importantly, unlike PE5, these additional PEs do not contain C-terminal Gly-Gly motifs (Figure 2C). When we performed directed co-IPs between KLHDC2 and these PEs, we found that most PEs could not interact with KLHDC2 (Figure 2D-E). However, some of PEs appear to be capable of interacting with KLHDC2 despite their lack of a cognate C-degron; PE20 interacted robustly with KLHDC2 despite ending in Ala-Val-Ser (Figure 2C-E). This demonstrates that KLHDC2 does not bind universally to PEs, but suggests that KLHDC2 may recognize additional C-degrons or that some PEs may be cleaved or processed in mammalian cells to reveal a KLHDC2-specific C-degron.

### PE5 is degraded after recognition by KLHDC2

Since PE5 is bound by the substrate binding domain of KLHDC2, we predicted that PE5 is a substrate of CRL2-KLHDC2. To establish this, we first measured the protein degradation of PE5 and PE5ΔGG using cycloheximide chase assays in RAW 264.7 cells ectopically expressing each. We found that while GFP-PE5ΔGG is stable over several hours, GFP-PE5 has a half-life of approximately 3 hr and is almost entirely degraded after 5 hr (Figure 3A-B). This difference in stability suggests that PE5’s ability to bind KLHDC2 might control its degradation. To further explore this, we used a variety of approaches to modulate the activity of KLHDC2 and examined steady-state GFP-PE5 levels under each condition. First, we treated cells with the pan-Cullin inhibitor MLN4924 or the proteasome inhibitor MG-132 and found that in MLN4924- and MG-132-treated cells, significantly more GFP-PE5 was expressed compared to untreated cells (Figure 3C-D). This was only true for PE5, not PE5ΔGG, which is not bound by KLHDC2. This indicates PE5 is degraded by the proteasome, and this requires CRL2 activity. Next, to specifically implicate KLHDC2 in PE5’s degradation, we used shRNAs to generate stable KLHDC2 knockdown RAW 264.7 cells (Figure S1B). KLHDC2 knockdown cells expressed significantly more PE5 than the control (Figure 3E-F). Here again, knockdown of KLHDC2 increased the stability of PE5 but not PE5ΔGG, indicating that the stability of PE5 is dependent on its binding to KLHDC2. To complement KLHDC2 knockdown experiments, we next measured the steady-state levels of PE5 in cells overexpressing KLHDC2. In cells expressing FLAG-KLHDC2, less PE5 was detected, but cells expressing FLAG-KLHDC2mut had PE5 levels similar to the control (Figure 3G-H). Moreover, the stability of PE5ΔGG was not affected by the overexpression of either wild-type or mutant KLHDC2 (Figure 3G-H). Together, these data suggest that PE5 is degraded by the proteasome after being bound and ubiquitinated by the CRL2-KLHDC2 complex.

**Figure 3.**
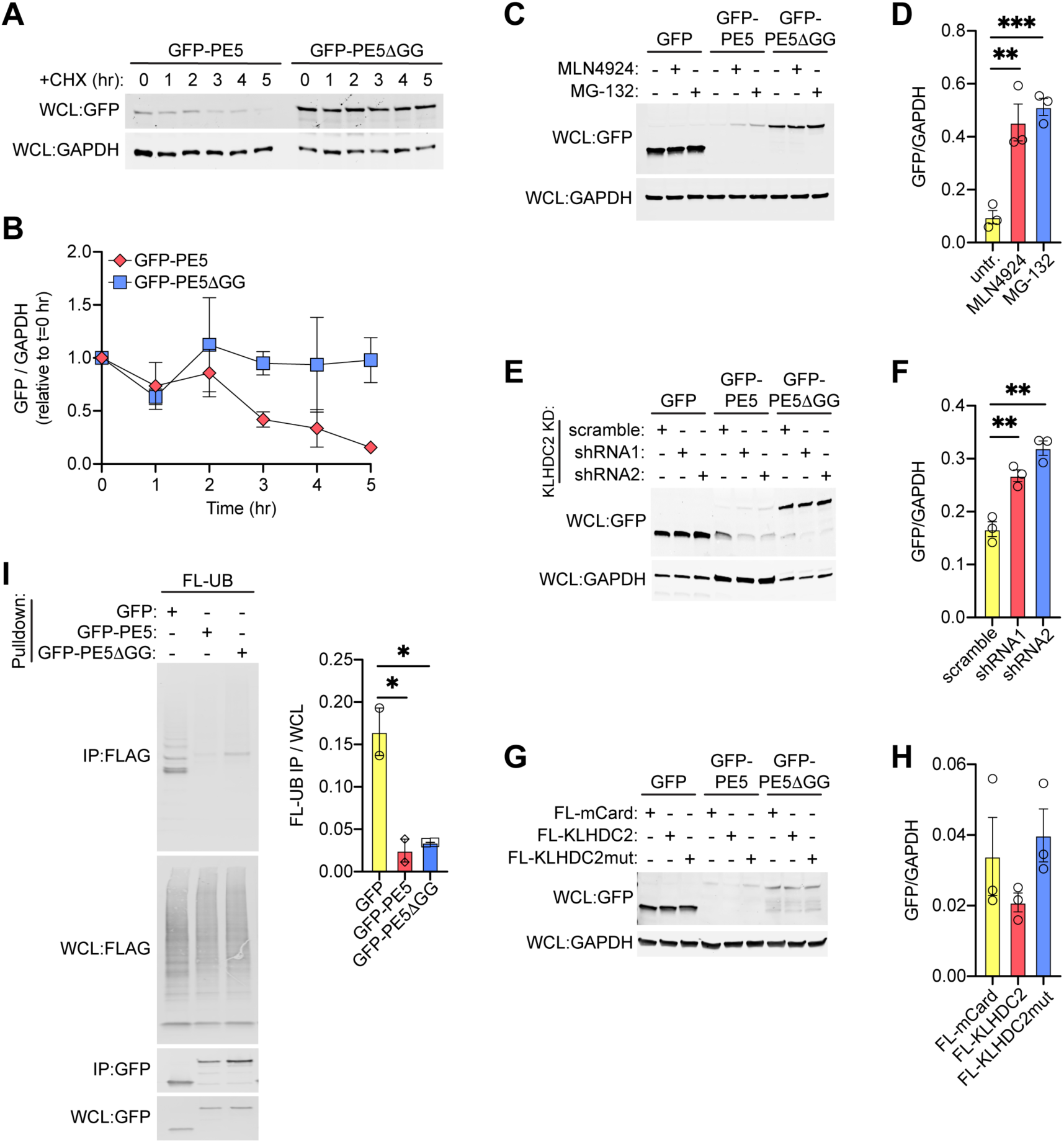
PE5 is degraded but not ubiquitinated by CRL2-KLHDC2. (A) Cell lysates from RAW 264.7 cells stably expressing GFP, GFP-PE5, or GFP-PE5ΔGG treated with cycloheximide (CHX; 5 mg/ml) for up to 5 hr. To detect PE5 at t=0 hr, cells were pre-treated with 5 mM MLN4924 and washed prior to addition of CHX. GFP probed to measure remaining PE5, and GAPDH probed as a loading control. (B) Quantification of A in which GFP is normalized to GAPDH, and the fraction of protein remaining at each time point is relative to t=0 hr. (C) Cell lysates from RAW 264.7 cells stably expressing GFP, GFP-PE5, or GFP-PE5ΔGG treated with MLN4924 (5 mM) or MG-132 (20 mM) as indicated for 2 hr. GFP probed to measure PE5 expression, and GAPDH probed as a loading control. (D) Quantification of C in which GFP signal is normalized to the GAPDH loading control. (E) Cell lysates from RAW 264.7 cells stably expressing shRNAs targeting KLHDC2 or a scramble shRNA as a negative control. Knockdown efficiency shown in Figure S1. GFP probed to measure PE5 expression, and GAPDH probed as a loading control. (F) Quantification of E in which GFP signal is normalized to the GAPDH loading control. (G) Cell lysates from RAW 264.7 cells stably expressing GFP, GFP-PE5, or GFP-PE5ΔGG along with 3xFLAG-tagged mCardinal (mCard), wild-type KLHDC2, or binding-deficient KLHDC2 S269E mutant (mut). GFP probed to measure PE5 expression, and GAPDH probed as a loading control. (H) Quantification of G in which GFP signal is normalized to the GAPDH loading control. (I) Denaturing IP (GFP-trap) of HEK293Ts transiently transfected with 3xFLAG (FL)-tagged Ubiquitin (UB) and GFP, GFP-PE5, or GFP-PE5ΔGG. Quantification of ubiquitination (right) expressed as the fold enrichment of FL-UB in the IP relative to the WCL. Results are representative of three independent experiments. *p < 0.05, **p < 0.005, ***p < 0.001 by two-tailed t-test Error bars indicate SD.

### PE5 is not ubiquitinated by CRL2-KLHDC2

We next measured the ubiquitination of PE5. Interestingly, though, PE5 does not contain any Lys residues, which are the canonical sites of ubiquitination (Figure S1A). However, noncanonical ubiquitination can occur at Ser, Thr, and Met residues. Therefore, we formally tested ubiquitination by co-expressing 3xFLAG-ubiquitin (UB) and GFP-PE5 in HEK293Ts, pulling down GFP-PE5, and probing for FLAG-UB. The IP of GFP exhibits multiple bands in a laddering pattern, indicative of polyubiquitination. Conversely, neither GFP-PE5 nor GFP-PE5ΔGG exhibit this banding or laddering pattern, demonstrating they are not polyubiquitinated (Figure 3I). This is consistent with the lack of Lys residues in both wild-type and mutant PE5.

Though not ubiquitinated, PE5 is degraded in a CRL2-KLHDC2-dependent manner (Figure 3A-H). This finding is contrary to a model in which PE5 is a substrate of CRL2-KLHDC2. Therefore, we hypothesized that PE5 may instead act as a competitive inhibitor of KLHDC2 and promote the degradation of this CRL2 substrate receptor. To test this, we first assessed the ubiquitination of KLHDC2 in the presence of PE5. We observed an increase in the ubiquitination of KLHDC2 when co-expressed with GFP-PE5, but not with GFP-PE5ΔGG (Figure 4A), suggesting PE5 promotes the autoubiquitination of KLHDC2. However, when we measured the steady-state protein levels of 3xFLAG-KLHDC2, co-expressing with GFP-PE5 in RAW 264.7 cells did not significantly change KLHDC2 levels (Figure 4B). This was also true for endogenous KLHDC2 levels, which were unchanged by the ectopic expression of GFP-PE5 (Figure 4C). Additionally, when we measured the protein degradation of either endogenous KLHDC2 or 3xFLAG-KLHDC2, we did not observe a change in the stability of KLHDC2 in the presence of GFP-PE5 (Figure 4D-E and S2). Together, these data indicate that although PE5 expression may enhance the autoubiquitination of KLHDC2, it is not sufficient to alter the protein levels of KLHDC2.

**Figure 4.**
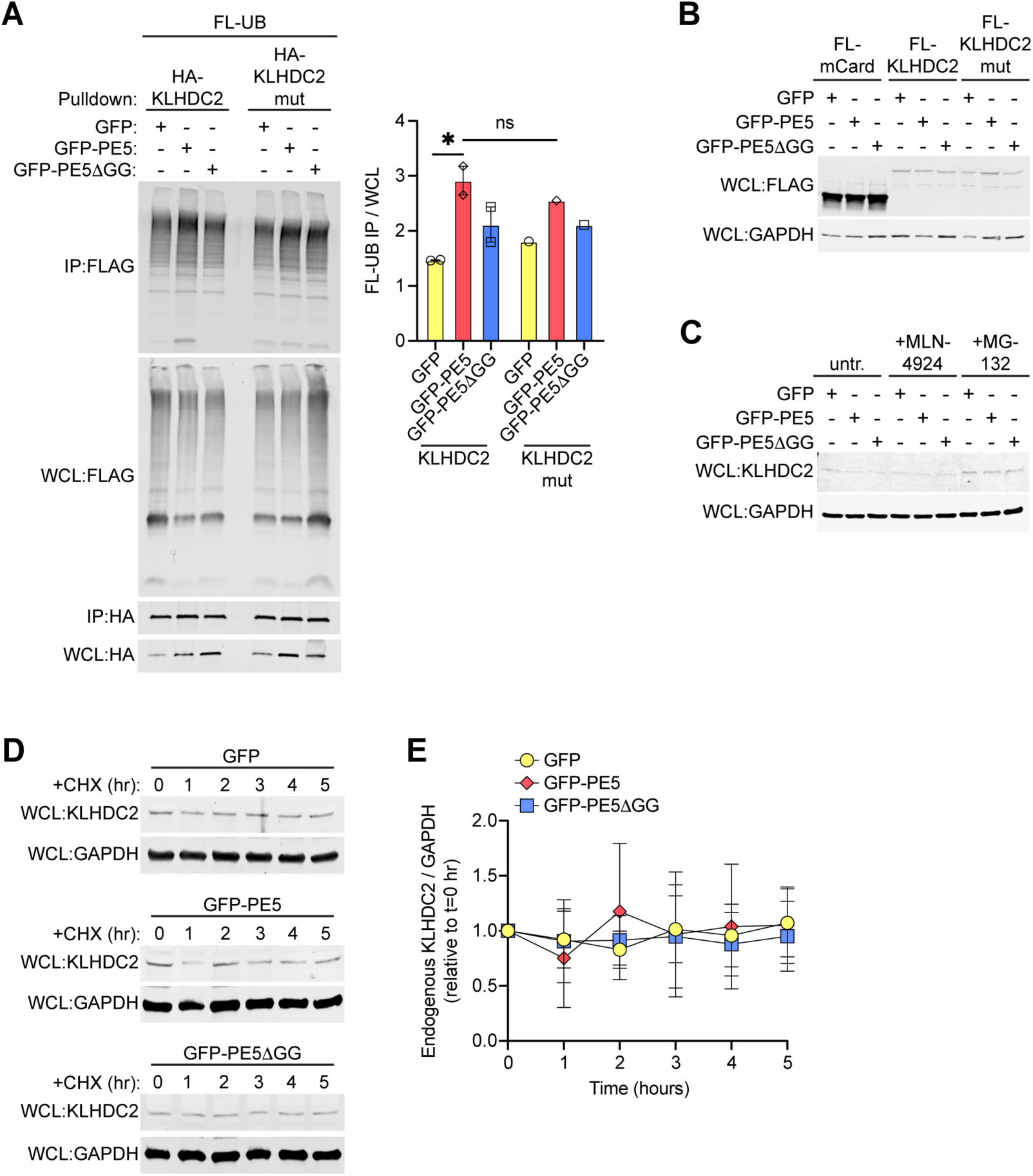
KLHDC2 is ubiquitinated but not degraded when PE5 is ectopically expressed. (A) Denaturing IP (HA resin) of HEK293Ts transiently transfected with 3xFLAG (FL)-tagged Ubiquitin (UB) with GFP, GFP-PE5, or GFP-PE5ΔGG and HA-tagged KLHDC2 or KLHDC2 S269E (mut). Quantification of ubiquitination (right) expressed as the fold enrichment of FL-UB in the IP relative to the WCL (n=2). (B) Cell lysates from RAW 264.7 cells stably expressing 3xFLAG-tagged mCardinal (mCard), wild-type KLHDC2, or binding-deficient KLHDC2 S269E mutant (mut) along with GFP, GFP-PE5, or GFP-PE5ΔGG. FLAG probed to measure KLHDC2 expression, and GAPDH probed as a loading control. (C) Cell lysates from RAW 264.7 cells stably expressing 3xFLAG-tagged mCardinal (mCard), wild-type KLHDC2, or binding-deficient KLHDC2 S269E mutant (mut) treated with MLN4924 (5 mM) or MG-132 (20 mM) as indicated for 2 hr. FLAG probed to measure KLHDC2 expression, and GAPDH probed as a loading control. (D) Cell lysates from RAW 264.7 cells stably expressing GFP, GFP-PE5, or GFP-PE5ΔGG treated with cycloheximide (CHX; 5 mg/ml) for up to 5 hr. Endogenous KLHDC2 probed to measure remaining protein, and GAPDH probed as a loading control. (E) Quantification of D in which KLHDC2 is normalized to GAPDH, and the fraction of KLHDC2 remaining at each time point is relative to t=0 hr (n=3). Results are representative of at least three independent experiments. *p < 0.05 by two-tailed t-test. Error bars indicate SD.

### PE5 does not change the degradation of CRL2 substrates

Since KLHDC2 stability is not altered by PE5, we next hypothesized that PE5 may alter the activity of CRL2-KLHDC2 or alternatively, the activity of other CRL2 complexes equipped with other substrate receptors. We first tested whether PE5 is an inhibitor of CRL2-KLHDC2 by measuring the degradation of the N-terminal domain (NTD) of USP1^17^, a known model substrate of CRL2-KLHDC2. We co-expressed 3xFLAG-NTD-USP1 and GFP-PE5 in RAW 264.7 macrophages and used cycloheximide chases to measure the degradation of NTD-USP1. We observed no difference in the stability or half-life of NTD-USP1 when expressed with wild-type or mutant PE5 (Figure 5A-B), indicating that PE5 does not prevent its degradation. Treatment with MLN4924 prevented degradation of NTD-USP1, verifying that it is a CRL2 substrate in RAW 264.7 cells (Figure S3A).

**Figure 5:**
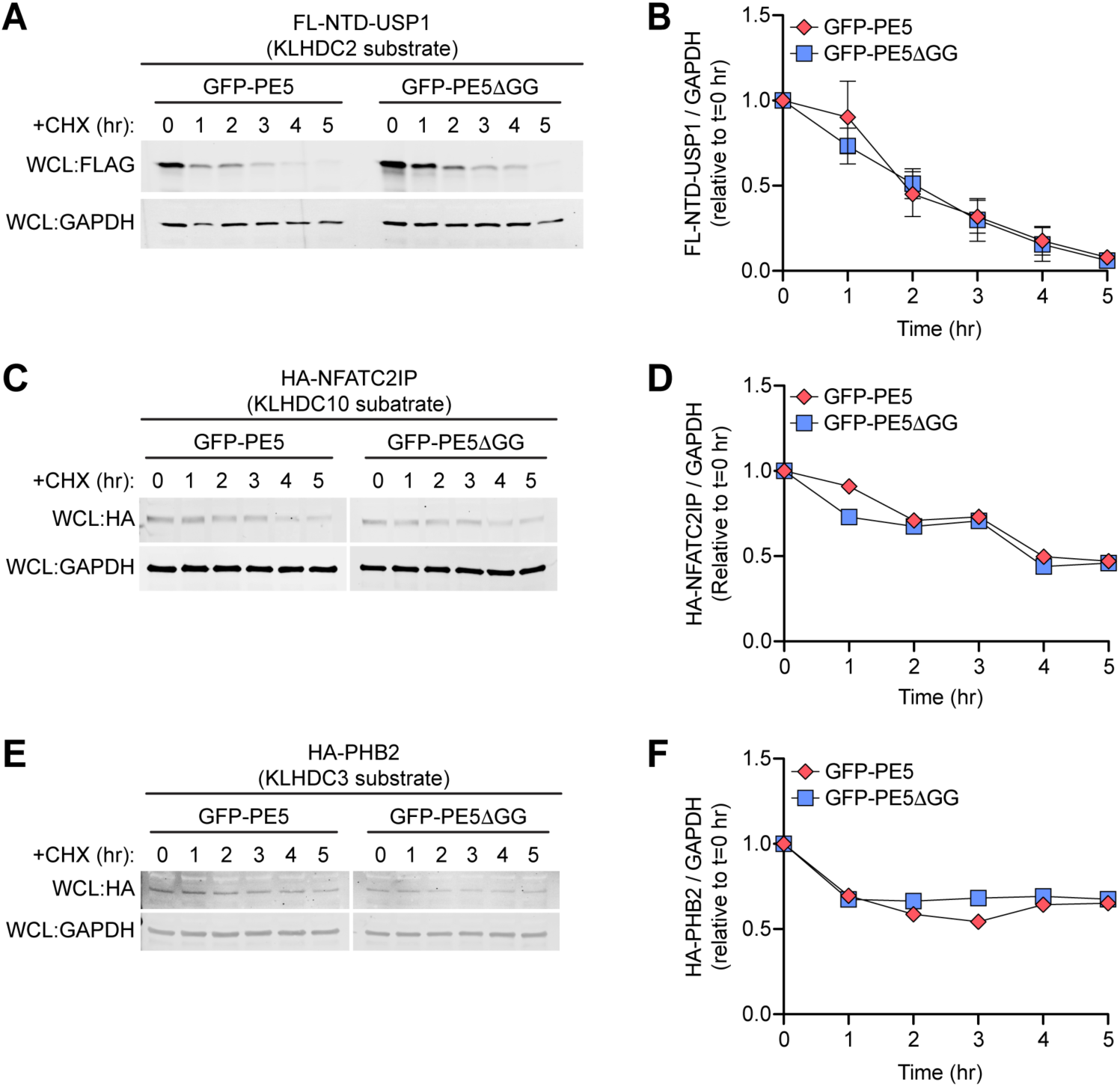
PE5 does not affect the degradation of model CRL2 substrates. (A) Cell lysates from RAW 264.7 cells stably expressing GFP-PE5 or GFP-PE5ΔGG and 3xFLAG-tagged NTD-USP1, a model substrate of KLHDC2, treated with cycloheximide (CHX; 5 mg/ml) for up to 5 hr. FL-NTD-USP1 probed to measure remaining protein, and GAPDH probed as a loading control. (B) Quantification of A in which FL-NTD-USP1 is normalized to GAPDH, and the fraction remaining at each time point is relative to t=0 hr (n=3). (C-D) As in A-B but for HA-tagged NFATC2IP, a model substrate of receptor KLHDC10. (E-F) As in A-B but for HA-tagged PHB2, a model substrate of receptor KLHDC3. Results are representative of at least three independent experiments. Error bars indicate SD.

We next tested whether PE5 might inhibit other CRL2 complexes. We co-expressed GFP-PE5 with either 3xFLAG-NFATC2IP, a substrate of CRL2-KLHDC10^26^, or 3xFLAG-PHB2, a substrate of CRL2-KLHDC3^25^. Similar to NTD-USP1, both NFATC2IP and PHB2 had similar stabilities and half-lives when expressed with either wild-type or mutant PE5 (Figure 5D-F), indicating PE5 does not affect their degradation. Here, too, treatment with MLN4924 prevented PHB2 degradation, demonstrating it too is a bona fide CRL2 substrate in macrophages (Figure S3B). Together, these data suggest that PE5 does not impede the activity of CRL2-KLHDC2 or other CRL2 complexes.

### PE5 promotes bacterial replication through KLHDC2

To investigate the functional implications of PE5’s interaction with KLHDC2 on bacterial pathogenesis, we used a model pathogen, *Salmonella enterica* Typhimurium, to measure how ectopic expression of PE5 affects macrophages’ cell-intrinsic antibacterial defenses. We pretreated PE5-expressing RAW 264.7 cells with MLN4924 to augment PE5 expression, infected them with *Salmonella,* and measured colony-forming units (CFUs) after 24 hr of infection. We found that *Salmonella* survived and replicated better in the presence of GFP-PE5 (Figure 6A), demonstrating that PE5 promotes bacterial infection. This increase in bacterial replication was not observed in cells expressing GFP-PE5ΔGG, indicating that the PE5-dependent enhancement of bacterial replication requires PE5’s interactions with KLHDC2 (Figure 6A). To further probe the role of KLHDC2 in controlling bacterial replication, we infected our KLHDC2 knockdown macrophages (Figure S1B) with *Salmonella* and measured survival and replication. Compared to control cells, KLHDC2 knockdowns failed to control *Salmonella* replication (Figure 6B). These data suggest that KLHDC2 is required for the host response to bacterial pathogens, making it a novel host restriction factor. Together, these data suggest that PE5’s ability to target this host factor may contribute to Mtb virulence and promote Mtb’s survival and replication in macrophages.

**Figure 6.**
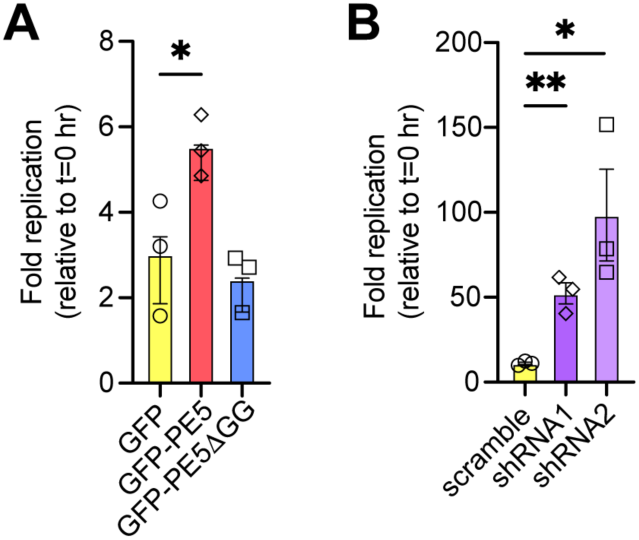
PE5 promotes bacterial replication through KLHDC2. (A) Salmonella replication in RAW 264.7 cells stably expressing GFP, GFP-PE5, or GFP-PE5ΔGG. Cells were infected with stationary phase Salmonella at a multiplicity of infection (MOI) of 10. Bacterial replication was assessed 24 hr post-infection and expressed as fold replication relative to t=0 hr (n=4). (B) Salmonella replication in KLHDC2 knockdown RAW 264.7 cells stably expressing shRNAs targeting KLHDC2 or a scramble shRNA as a negative control. Cells were infected and Salmonella replication was quantified as in A (n=4). Results are representative of three independent experiments. *p < 0.05, **p < 0.005 by two-tailed t-test. Error bars indicate SD.

## DISCUSSION

Here we show that PE5 interacts with the host ubiquitin ligase complex, CRL2. It binds to this complex via the substrate receptor KLHDC2, which recognizes the Gly-Gly C-degron in PE5. Binding to PE5 is specific to KLHDC2, as other substrate receptors are incapable of recognizing PE5. Additionally, KLHDC2 does not recognize all PE/PPE proteins, as its binding to PE5 is specific and depends on PE5’s C-terminal Gly-Gly motif. While PE5 is not itself ubiquitinated by the CRL2 complex that recognizes it, PE5 is still degraded in a CRL2-KLHDC2-dependent manner. The mechanism of this is still unclear, but it suggests that PE5 is likely delivered to the proteasome by a ubiquitinated binding partner. While we initially suspected that PE5’s ubiquitinated binding partner was KLHDC2, KLHDC2 is not degraded in the presence of PE5. Interestingly, PE5 does not appear to alter the activity CRL2-KLHDC2 or other CRL2 complexes, indicating that PE5 does not inhibit the function of this ubiquitin ligase complex. Nonetheless, ectopic expression of PE5 in macrophages promotes intracellular bacterial infection, and knockdown of its target, KLHDC2, similarly enhances replication of intracellular bacteria. Together, these data indicate that PE5 is an Mtb virulence factor that targets a host ubiquitin ligase critical for cell-intrinsic responses to infection.

Despite identifying a host target of PE5 and demonstrating that ectopic expression of PE5 promotes intracellular bacterial replication, it remains unclear how precisely PE5 functions through CRL2-KLHDC2 to block cell-intrinsic defenses. Our initial hypothesis was that PE5 blocks CRL2 activity by acting as an unproductive substrate that cannot be ubiquitinated. We predicted that this could block CRL2-KLHDC2 function or prevent the assembly of other CRL2 complexes. However, our data indicate that the activity of these complexes does not decrease when PE5 is ectopically expressed. An alternative possibility is that PE5 instead acts as a carrier to deliver non-typical substrates to CRL2-KLHDC2. In this way, PE5 could enhance the ubiquitination and degradation of host factors critical for responding to bacterial infection. Indeed, hijacking CRL2 complexes like this has been observed for at least one viral effector. The herpesvirus protein UL49.5 binds to TAP (transporter associated with antigen presentation), and delivers it to CRL2-KLHDC3, promoting its ubiquitination and degradation, and ultimately inhibiting antigen presentation during viral infection^25^. It’s tempting to speculate that PE5 similarly drags host factors, such as trafficking proteins, lysosomal proteins, transcription factors, signaling proteins, etc., to CRL2-KLHDC2 to force their ubiquitination and degradation. While our AP-MS datasets do not contain any obvious PE5 binding partners that would fit this model, it is possible that PE5 expression rapidly degrades its partner, making it difficult to detect in standard AP-MS. Additionally, PE5 may capture partners that are only present in macrophages; since our AP-MS was performed in non-immune HEK293Ts, it is possible they are absent from our AP-MS data due to cell type specificity. Our ongoing work is determining what host proteins may be destabilized as a result of PE5 potentially hijacking CRL2-KLHDC2.

While we did not identify any PE5-dependent stabilization of known CRL2 substrates, it is possible that in macrophages, CRL2 complexes target a unique repertoire of substrates that has yet to be fully defined. Indeed, the cataloging of CRL2 substrates and the identification of well-defined C-degrons for each receptor is an active field of study. It would not be surprising if CRL2 complexes have cell-type-specific substrates, but most work on these complexes has been performed in non-immune cells. Future studies will seek to define the repertoire of substrates regulated by each CRL2 complex and how PE/PPEs like PE5 might affect their stability both at rest and in response to infection. While CRL2 complexes have been implicated in cellular processes like cell death pathways, cellular stress responses, cell proliferation, and more, their regulation in immune cells and the role they play during infection have not been deeply explored. However, our data indicate that KLHDC2 is required for controlling *Salmonella* infection in macrophages, indicating it functions as a host restriction factor. However, it’s unknown if the altered stability of KLHDC2 substrates is key for this antibacterial function, and if so, what substrates contribute to this host restriction. Alternatively, KLHDC2 could function in an unknown independent manner to promote antibacterial defenses.

Past studies on PE5 have focused on its function in iron acquisition^13,27^. However, these studies have not established a defined role for PE5 in this process. PE5 forms a heterodimer with PPE4 to facilitate secretion through the ESX-3 secretion system. PPE4 is not detected in culture supernatants, suggesting it may remain on the surface of Mtb while PE5 is released into the bacterium’s surroundings^9–11^. Surface retention of PPE4 would be consistent with the predicted and demonstrated localization of many other PPE proteins and may suggest a role for PPE4 in forming a channel or exporter in the mycomembrane. Conversely, PE5’s release from the bacterium would be consistent with PE5 acting as a bona fide virulence factor capable of directly interfacing with host cell biology. However, because deletion of PE5 and PPE4, which are essential, is possible only when iron is supplemented, testing the specific contribution of PE5 to host-pathogen interactions, where iron remains sufficiently limited to prevent growth of a PE5 mutant, has been an insurmountable challenge. By identifying a mutant of PE5, PE5ΔGG, we have introduced a novel opportunity to better probe the effector function of PE5 inside a host cell. The small C-terminal deletion in PE5ΔGG does not affect the PE domain required for PE5’s interaction with PPE4, and therefore, it is not expected to affect their secretion or the function of PPE4. However, our work has established that PE5ΔGG is incapable of binding to CRL2-KLHDC2 and getting degraded in the host cell, making it a valuable genetic tool for studying this PE protein during Mtb infection. Future work will use this PE5ΔGG variant to probe, for the first time, PE5’s contribution to Mtb growth and survival in macrophage and mouse infection models.

Our work identified a novel interaction between PE5’s C-degron and one C-degron receptor. While no other PE proteins contain the Gly-Gly C-degron recognized by KLHDC2, some PPE proteins contain this C-degron, and several other PE proteins contain other substrate receptors’ C-degrons. This raises the possibility that binding to CRL2 complexes may be a broader strategy used by Mtb to either block the host’s CRL2 complexes or hijack CRL2 complexes to degrade host restriction factors. However, containing a C-degron is not sufficient for binding to substrate receptors; the C-terminus must be exposed and flexible enough to enter the deep binding pocket of the substrate receptors. Additional work is needed to systematically assess the interactions between PE/PPE proteins and CRL2 receptors to determine whether this is a unique functionality of PE5 toward CRL2-KLHDC2 or if many mycobacterial PE/PPE proteins can target a variety of CRL2 complexes to block antibacterial defenses.

## MATERIALS & METHODS

### Cell lines and cell culture

RAW 264.7 cells (ATCC TIB-71), HEK293T cells (ATCC CRL-3216), and Lenti-X 293T cells (TaKaRa Bio) were cultured in Dulbecco’s Modified Eagle Medium (DMEM) supplemented with 10% heat-inactivated fetal bovine serum (FBS) and 20 mM HEPES, at 37℃ with 5% CO2. When required, RAW 264.7 cells were selected and maintained in 5 µg/ml puromycin or 10 μg/ml blasticidin. For protein analysis, cycloheximide chases, and infections, cells were plated at 5 x 10^5^ cells/well in a 12-well tissue culture-treated (TC) dish. For infections, selection antibiotics were omitted from culture media.

### Molecular cloning

PE/PPEs were cloned from *M. tuberculosis* H37Rv genomic DNA and CRL2 proteins from cDNA from RAW 264.7 cells. Genes were inserted via Gibson assembly (NEB) into pENTR1a entry vectors modified to contain in-frame HA-, 3xFLAG- or GFP-tags, as previously described (REF – gal8). Mutations were introduced by traditional cloning (PE5DGG) or by site-directed mutagenesis with primers designed using PrimerX (https://www.bioinformatics.org/primerx/). All constructs were fully Sanger sequenced (Azenta) to ensure in-frame, error-free tagged proteins. Sequenced constructs were then gateway cloned using LR Clonase (Invitrogen) into pLenti destination vectors (Addgene, plasmid 19067)^28^. Expression of tagged proteins was validated in HEK293T cells via transient transfection with 2 μg DNA and PolyJet (SignaGen). After 1 to 2 days, cells were harvested and proteins were separated by SDS-PAGE and visualized by Western blot using primary antibodies specific to FLAG (clone M2; Sigma-Aldrich, catalog no. F1804), HA (Roche, catalog no. 11867423001), GFP (Abcam, catalog no. ab183734), or KLHDC2 (Invitrogen, catalog no. PIPA590252), Li-Cor 700 and 800 secondary antibodies, and the Li-Cor Odyssey Fc imager.

Oligonucleotides for shRNAs targeting KLHDC2 were designed using the Designer of small interfering RNA website (http://biodev.cea.fr/DSIR/DSIR.php) and synthesized by Sigma. Oligos were annealed and ligated into HpaI- and XhoI-digested pSicoR-GFP. Constructs were confirmed by restriction digest and Sanger sequencing to ensure insertion shRNAs. Knockdowns were validated by RNA isolation (Zymo kits) and RT-qPCR (Bio-Rad iScript, Quanta SYBR Fast, Bio-Rad 384 CFX) using KLHDC2 and Actin primers.

For stable expression in RAW 264.7 cells, lentiviral transduction was used. Lentivirus was produced by co-transfecting Lenti-X 293T cells with lentiviral plasmids and the packaging plasmids psPAX2 and pMD2G/VSV-G (Addgene, plasmids 12259 and 12260). RAW 264.7 cells were transduced with lentivirus using Lipofectamine 2000 (Invitrogen, diluted 1:1000) and selected with appropriate antibiotics for 3 to 5 days. Stable expression was confirmed by Western blot using an epitope-specific primary antibody^28^.

### Immunoprecipitations

RAW 264.7 stable expression cells were plated in 6-cm TC dishes at 80% confluency for next-day IPs. HEK293Ts were plated in a 6-cm TC dishes at 40% confluency for next-day transfection prior to IPs. HEK293Ts were transfected with 4 μg of expression plasmid (2 μg of each expression plasmid to test protein-protein interactions) using PolyJet. After 24 to 48 hours, cells were washed and lifted in PBS + 4 mM EDTA, pelleted by centrifugation at 1,000 x g for 5 min. Cells were lysed in lysis buffer (150 mM Tris HCl pH 7.5, 50 mM NaCl, 1 mM EDTA, 0.075% NP-40, protease inhibitors [Roche]). Cell lysate was cleared of cellular debris and nucleic acids by centrifugation at 7,000 x g for 10 min. 5% of the cleared cell lysate was boiled with 4x Laemmli sample buffer (Bio-Rad) and 2-mercaptoethanol (Bio-Rad) and saved as the “whole cell lysate”. The remaining cleared cell lysate was incubated with a pre-washed (three times in lysis buffer) 20 μl GFP trap (Proteintech cat. no. gta) or antibody-conjugated beads/resin (FLAG [EZview Red anti-FLAG M2 affinity gel, Sigma-Aldrich]; HA [Pierce anti-HA agarose, Thermo Scientific]) for 30 to 60 min at 4°C with rotation. Beads were washed three times with wash buffer (150 mM Tris pH 7.5, 50 mM NaCl, 1 mM EDTA, 0.05% NP-40). Proteins were eluted either by boiling for GFP-tagged proteins or with an excess of competing peptide (3xFLAG peptide [Sigma] for FLAG-tagged proteins or HA peptide [Sigma] for HA-tagged proteins) resuspended in wash buffer. Eluates were boiled with 4x Laemmli sample buffer and 2-mercaptoethanol, and protein-protein interactions were visualized by Western blot using epitope-specific antibodies^20,28^.

### Mass spectrometry

To identify novel protein-protein interactions, co-IP eluates were run on SDS-PAGE gel. The gel was fixed in 50% methanol and 10% acetic acid for 1 hour, stained with Coomassie blue (Bio-Rad) for 1 hour, and destained in ultrapure water overnight. Mass spectrometry analysis was performed by the Rutgers Center for Advanced Proteomics Research (CAPR). In-gel trypsin digestion was performed, and peptides were desalted and analyzed by LC-MS/MS on Orbitrap Fusion Lumos mass spectrometer coupled with an Ultimate 3000 nano HPLC (Thermo Scientific). Peptides were identified by searching MS/MS spectra against the Uniprot human database using Spectronaut software.

### Protein stability assays

Whole cell lysate was prepared from confluent cells grown in 6-well TC dishes and lysed in 400 µl of lysis buffer (1% NP40, made in 50 mM HEPES, pH 7.4 and 150 mM NaCl). Lysates were spun at 7,000 x g for 5 min to remove nuclei. 4x LDS sample buffer and 2-mercaptoethanol were added to each sample, and samples were boiled at 95°C for 5 min. Lysates were run on Any-KD SDS-PAGE gels (Bio-Rad) and electrophoresis was run at 100 V for approximately 1.5 hr in the Mini-PROTEAN System (Bio-Rad) in running buffer (25 mM Tris, 192 mM Glycine, 0.1% SDS). Separated proteins were transferred onto nitrocellulose membrane (Bio-Rad) in the same Mini-PROTEAN System at 100 V for 1 hr in transfer buffer (25mM Tris pH 8.3, 192 mM Glycine, 20% methanol). The membrane was blocked for 30 minutes in Intercept PBS Blocking Buffer (Licor) and incubated with primary antibody on a rocker overnight at room temperature. Primary antibodies were diluted in Intercept PBS blocking buffer (Licor): 3xFLAG (1:5000, Sigma, catalog no. F3165), HA (1:5000, Roche, catalog no. 11867423001), GFP (1:5000, Abcam, catalog no. ab183734), and KLHDC2 (1:1000, ThermoFisher, catalog no. PA5-90252). Membranes were washed three times in TBST (TBS, 0.1% Tween) for 10 minutes and incubated with fluorophore-conjugated (680 nm and 800 nm) secondary antibodies diluted 1:20,000 in blocking buffer and incubated at room temperature for 1.5 hr. Membranes were washed 3 times in TBST, once in TBS, and imaged on a LiCor Odyssey Fc imaging system.

### Bacterial infections

RAW 264.7 cells were plated at 5 x 10^5^ cells/well in a 12-well TC dish using 4 biological replicates per condition per time point. The following day, an overnight culture of *Salmonella enterica* Typhimurium (grown in Luria-Bertani (LB) broth supplemented with 0.3 M NaCl) was washed twice in Hanks’ Balanced Salt Solution (HBSS). OD600 was used to determine the volume of bacteria needed for a multiplicity of infection (MOI) of 10 (1 OD600 = 2 x 10^9^ bacteria/ml). Bacteria were diluted in HBSS and added to RAW 264.7 cells. Infections were synchronized by spinning for 10 min at 1,000 x g, and cells were incubated at 37℃ with 5% CO_2_ for 20 min. Following infection, cells were washed with HBSS containing 100 μg/ml gentamicin and then cultured in media containing 5 μg/ml gentamicin for 24 hours. At 0 and 24 hours, cells were washed twice with PBS and lysed in 0.5% TritonX-100. Lysates were serially diluted 1:10 in PBS and spread on LB agar plates. Plates were incubated at 37°C overnight before enumerating colonies^28^.

## ACKNOWLEDGEMENTS

Mass spectrometry analysis was performed by Drs. Tong Liu and Hong Li at the Rutgers NJMS Center for Advanced Proteomics Research, which is partially supported by NIH grant 1S10OD034300 to H.L.. Select models and graphics were generated using BioRender (www.biorender.com) as noted in figure legends. We thank the members of the Bell Lab, past and present, for their help and thoughtful feedback. This work was supported by NIH grant DP2AI154429 to S.L.B., American Heart Association Postdoctoral Fellowship 25POST1365111 to B.T.S.A.M., and American Heart Association Predoctoral Fellowship 24PRE1198252 to J.R..

## AUTHOR CONTRIBUTIONS

Conceptualization – B.T.S.A.M., S.L.B.; Investigation – B.T.S.A.M., O.V., A.H., J.R. C.R., S.L.B.; Data Analysis - B.T.S.A.M., O.V., A.H., J.R. C.R., S.L.B.; Writing – B.T.S.A.M., S.L.B.; Visualization – B.T.S.A.M., S.L.B.; Funding acquisition – S.L.B., B.T.S.A.M, J.R.; Supervision – S.L.B.

---

**Figure S1.**
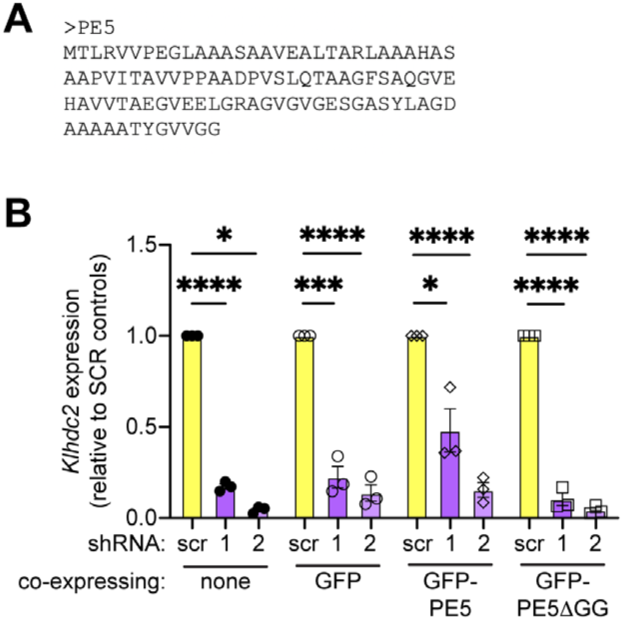
(A) Protein sequence of PE5, which includes no Lys residues. (B) RT-qPCR analysis of Klhdc2 in RAW 264.7 cells carrying shRNAs targeting KLHDC2 (two independent shRNA constructs #1 and #2) or a negative control scramble shRNA. Knockdowns were also generated independently in RAW 264.7 cells expressing GFP, GFP-PE5, or GFP-PE5DGG. Klhdc2 mRNA levels normalized to Actb levels and calculated as a ratio relative to the scramble shRNA for each parental cell line. p < 0.05, **p < 0.005, ***p < 0.001 by two-tailed t-test Error bars indicate SD.

**Figure S2.**
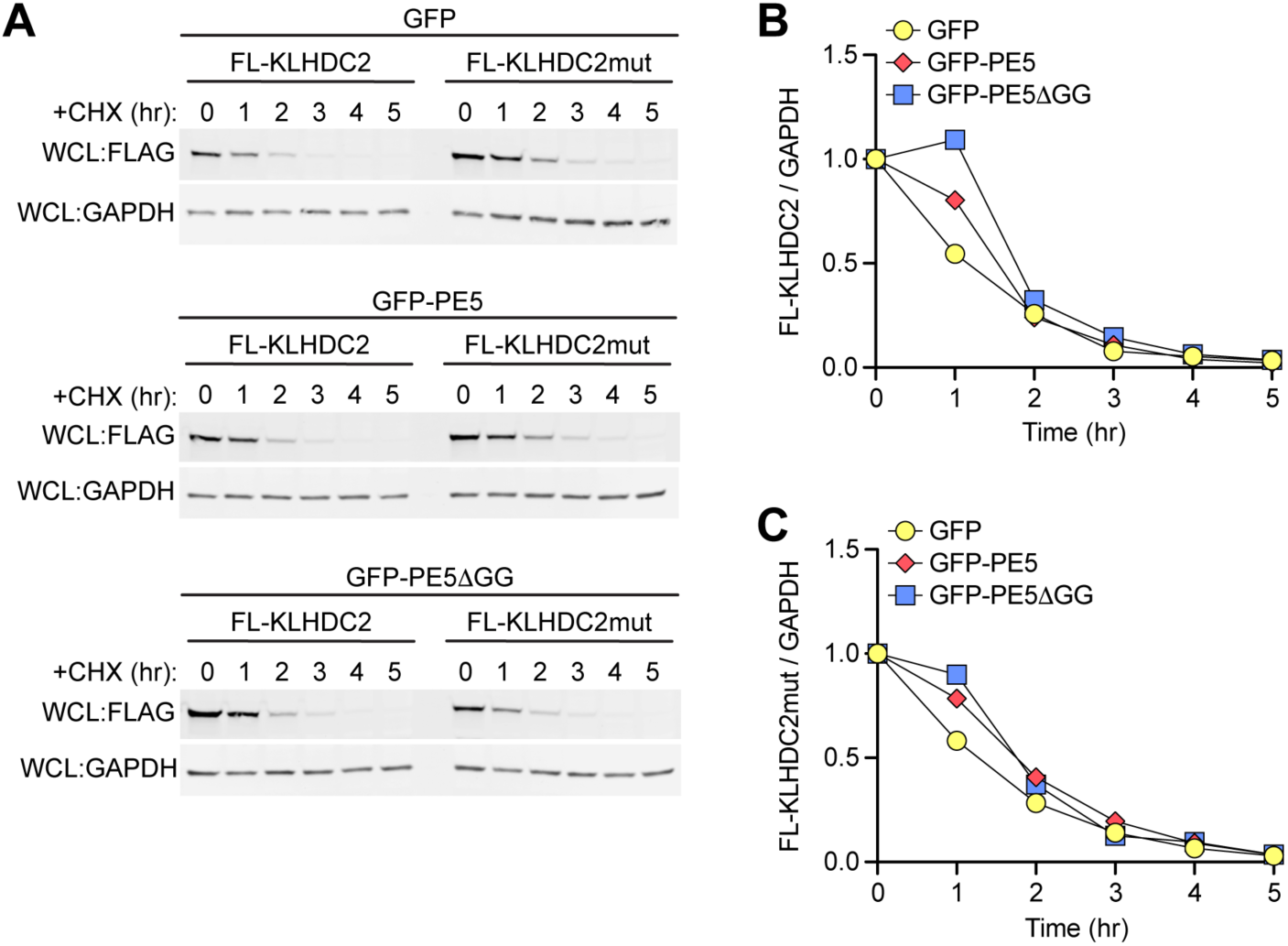
(A) Cell lysates from RAW 264.7 cells stably expressing GFP, GFP-PE5, or GFP-PE5ΔGG and 3xFLAG-tagged KLHDC2 or KLHDC2 S269E (mut) treated with cycloheximide (CHX; 5 mg/ml) for up to 5 hr. FLAG probed to measure remaining FL-KLHDC2, and GAPDH probed as a loading control. (B) Quantification of A in which FL-KLHDC2 is normalized to GAPDH, and the fraction remaining at each time point is relative to t=0 hr. (C) As in B but for FL-KLHDC2mut.

**Figure S3.**
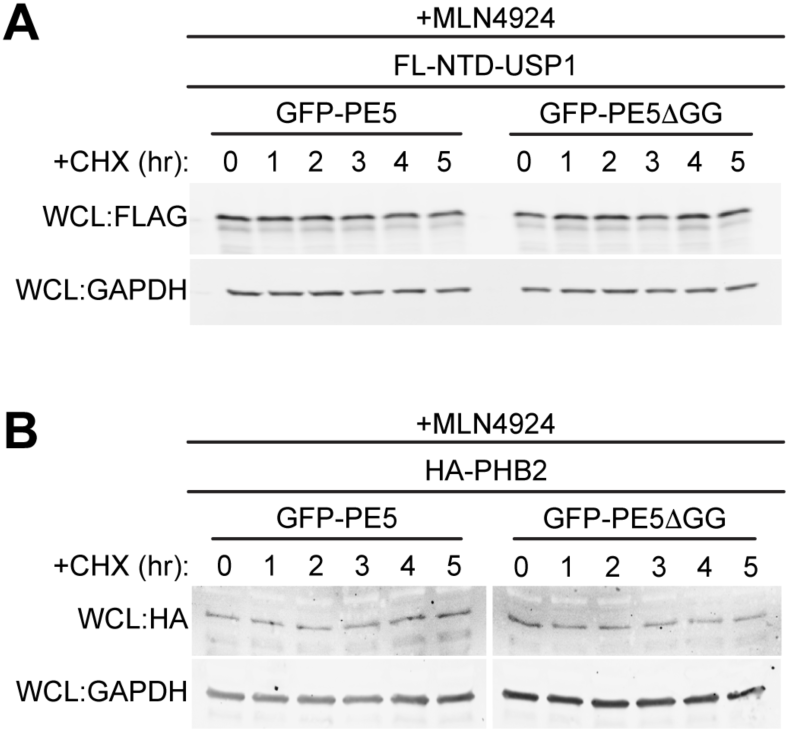
(A) Cell lysates from RAW 264.7 cells stably expressing GFP-PE5 or GFP-PE5ΔGG and 3xFLAG-tagged NTD-USP1, a model substrate of KLHDC2, treated with cycloheximide (CHX; 5 mg/ml) and MLN4924 (5 mM) for up to 5 hr. FL-NTD-USP1 probed to measure remaining protein, and GAPDH probed as a loading control. (B) As in A but for HA-tagged PHB2, a model substrate of KLHDC3. Results are representative of three independent experiments.

